# Enhanced cell segmentation with limited annotated data using generative adversarial networks

**DOI:** 10.1101/2023.07.26.550715

**Authors:** Abolfazl Zargari, Najmeh Mashhadi, S. Ali Shariati

## Abstract

The application of deep learning is rapidly transforming the field of bioimage analysis. While deep learning has shown great promise in complex microscopy tasks such as single-cell segmentation, the development of generalizable foundation deep learning segmentation models is hampered by the scarcity of large and diverse annotated datasets of cell images for training purposes. Generative Adversarial Networks (GANs) can generate realistic images that can potentially be easily used to train deep learning models without the generation of large manually annotated microscopy images. Here, we propose a customized CycleGAN architecture to train an enhanced cell segmentation model with limited annotated cell images, effectively addressing the challenge of paucity of annotated data in microscopy imaging. Our customized CycleGAN model can generate realistic synthetic images of cells with morphological details and nuances very similar to that of real images. This method not only increases the variability seen during training but also enhances the authenticity of synthetic samples, thereby enhancing the overall predictive accuracy and robustness of the cell segmentation model. Our experimental results show that our CycleGAN-based method significantly improves the performance of the segmentation model compared to conventional training techniques. Interestingly, we demonstrate that our model can extrapolate its knowledge by synthesizing imaging scenarios that were not seen during the training process. Our proposed customized CycleGAN method will accelerate the development of foundation models for cell segmentation in microscopy images.

## Introduction

Generative Adversarial Networks (GANs) have gained significant attention in recent years due to their remarkable success in generating realistic images and videos [1, 2]. GANs are deep learning architecture that consists of two neural networks: a generator and a discriminator [3-5]. The generator network is responsible for synthesizing new data, while the discriminator network attempts to distinguish between real and synthetic samples. The two networks compete in a game-like scenario until the generator produces data that is nearly indistinguishable from real data. Generative Adversarial Networks (GANs) have exhibited substantial capabilities across a wide variety of applications, including image synthesis [6], video generation [7], and natural language processing [8].

Cell segmentation is a crucial step in microscopy, which involves the identification and delineation of individual cells within images [9-11]. Cell segmentation is inherently complex due to the diversity and irregularity of morphological features of different cell types, such as shape and size, as well as the propensity for cells to cluster together, making highly accurate segmentation a challenging task. Deep learning methods, particularly convolutional neural networks (CNNs), have shown great success in improving cell segmentation accuracy [12-14]. The development and application of deep learning models for cell segmentation heavily depend on the availability of a large amount of annotated training data [15].

A significant bottleneck in the field of cell segmentation is the lack of sufficient annotated cell image datasets [16]. Manual annotation of cell images, which involves outlining individual cells in microscopic images to create “ground truth” masks, is a laborious and time-consuming task that requires considerable effort by cell biology experts[17-19]. It becomes practically infeasible when dealing with large volumes of data or when timely results are needed. This scarcity of annotated data impedes the training of robust models, limiting the potential improvements in cell segmentation tasks.

In this study, we propose a solution to tackle this challenge by employing a customized CycleGAN-based model to train a cell segmentation model with limited annotated data. We aimed to utilize the power of GANs to generate realistic and diverse cell images, thereby enriching the training data for the segmentation model. We systematically compared the CycleGAN-based training approach with that of the conventional training approach on the performance of our segmentation model. Our results showed that the CycleGAN-based approach, with limited annotated cell image datasets, increased the model’s ability to generalize and thus potentially enhancing the performance of cell segmentation tasks.

## Results

### Designing and training the segmentation model

An overview of our approach is illustrated in Figure 1A. The architecture introduced in this study uses a new modified CycleGAN approach for cell image segmentation, offering enhanced data diversity. The new features include a style generation path within a 2D-UNET-based image generator, two adversarial PathGan discriminators enhanced with a linear attention layer, and a balanced utilization of different loss functions. It also leverages a differentiable image augmentation technique, collectively resulting in realistic synthetic cell images, accurate segmentation, and improved training stability, especially with limited data.

**Figure 1.**
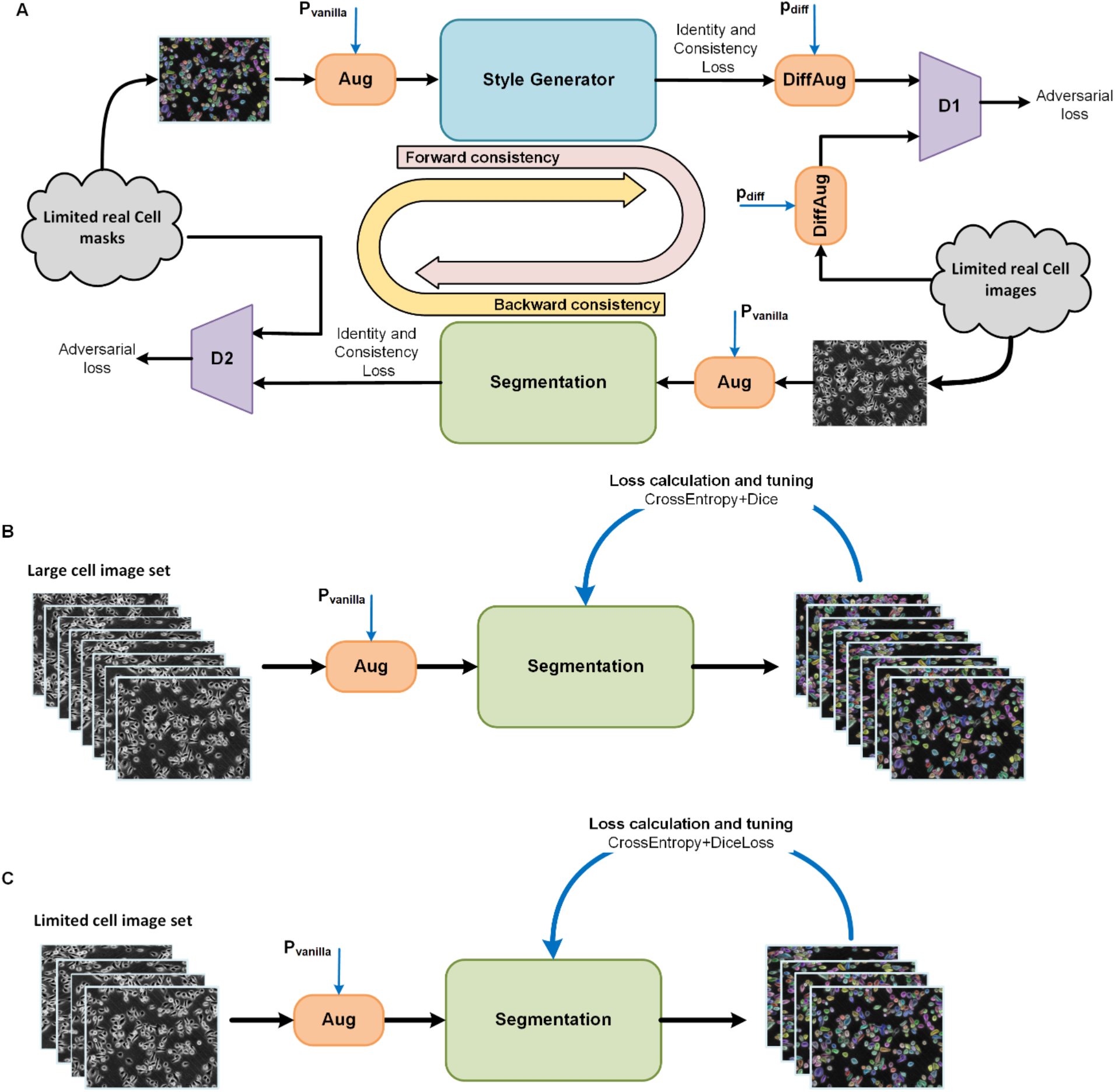
Three different cell segmentation training scenarios. A) Training the segmentation model in a CycleGAN-based process on a limited dataset of cell images involving the style generator and differentiable augmentation techniques. B) Training the segmentation model on a large dataset of cell images and applying vanilla augmentation pipeline (Aug). C) Training the segmentation model on a limited dataset of cell images (e.g., 25% original large dataset in this paper).

To test our proposed CycleGAN segmentation training method, we used our recently published annotated dataset of phase-contrast images of the DeepSea model [12] to compare three segmentation training scenarios: 1) Training the segmentation model on a smaller subset (25%) of full DeepSea training dataset (representing a selected limited dataset) involving a Cycle-GAN-based augmentation process and testing it on the DeepSea test set (Figure 1A), 2) Training the segmentation model on the 100% full DeepSea training dataset and testing it on the DeepSea test set (Figure 1B), and 3) Training the segmentation model on a smaller subset (25%) of full DeepSea training dataset and testing it on the same DeepSea test set (Figure 1C). The DeepSea dataset consists of a collection of annotated phase contrast time-lapse microscopy images that are difficult to segment because of low contrast, noisy, hard, and touching cell samples [12]. We used phase-contrast images of two cell type samples (Embryonic Stem Cells and Bronchial Epithelial Cells) from this dataset to demonstrate the versatility of our proposed method, underlining its applicability to various cell datasets with variations in cell shapes and sizes. We used the DeepSea baseline architecture as the segmentation model, which is an efficient scaled-down version of UNET [12]. In all three scenarios, during the training process, we passed the original training cell images and their corresponding binary mask images to a pipeline of a set of diverse conventional image augmentation functions in a randomized order (Figure S1A). The probability p_vanilla (taking any random value between 0 and 1) controls the frequency of augmentation requests during the training process. The augmentation process can help the model see more data variations during training and, thus, increase the model generalization while reducing the risk of overfitting [20]. However, it might yield very modest benefits when the training dataset is limited [21].

**Figure 2.**
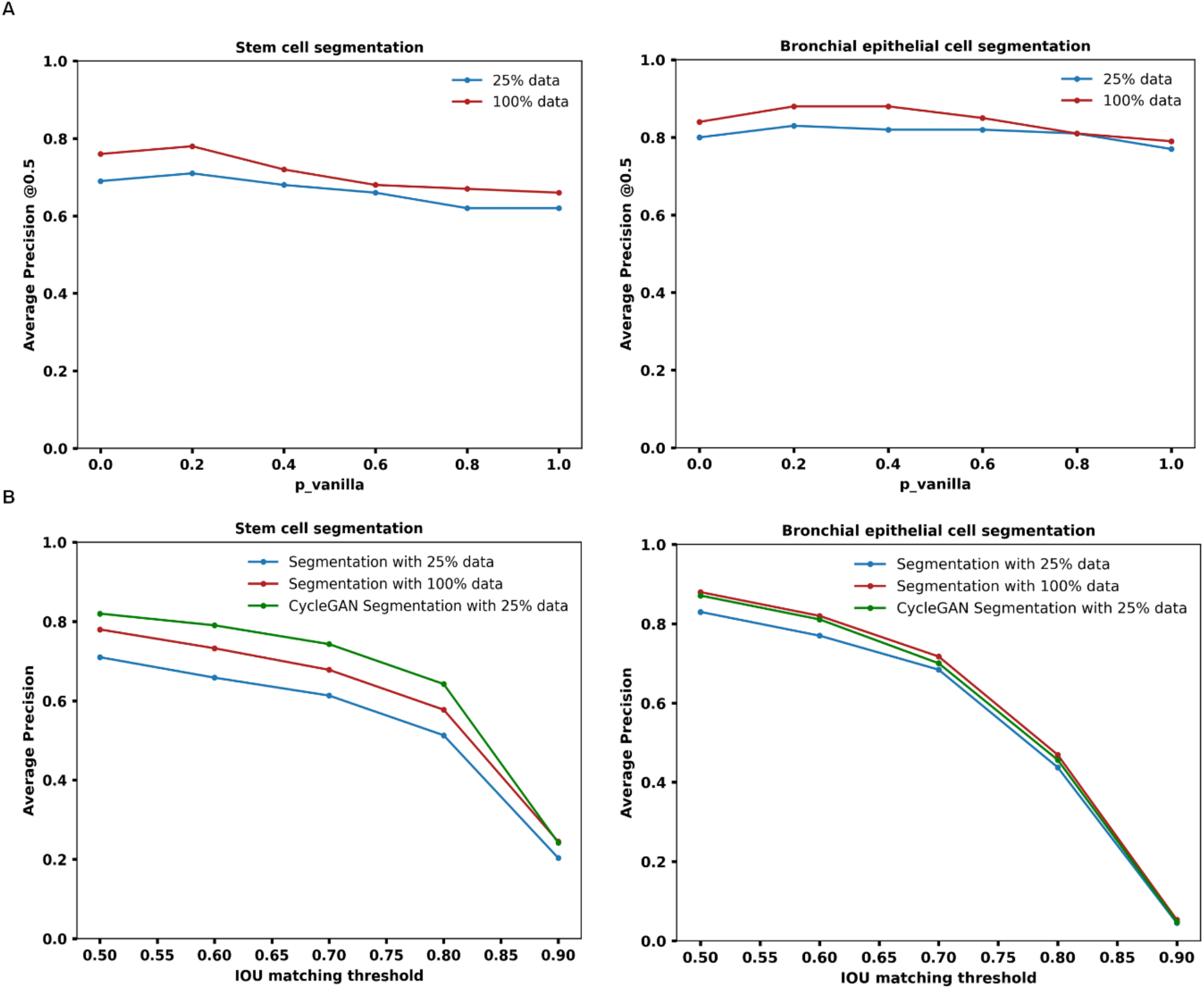
Enhanced cell segmentation performance by proposed CycleGAN-based training method. A) Vanilla augmentation performance effect on segmentation precision achieved with different training data sizes. B) The proposed CycleGAN-based training method compared to the segmentation model trained with 25% and 100% of the training dataset.

**Figure 3.**
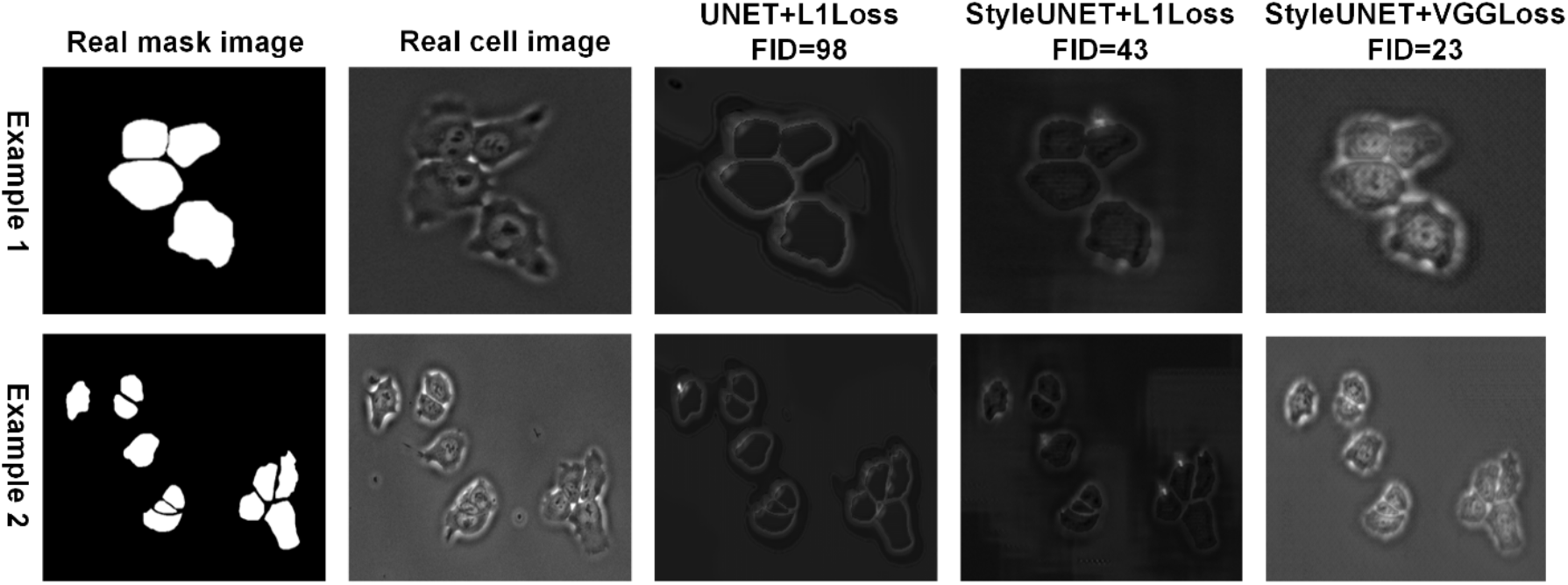
Improved realism of synthetic images generated by our modified CycleGAN. Two examples of the effect of the Style injecting technique and Vgg perceptual feature loss function on addition of morphological details and nuances to synthetic images.

In our proposed method (Figure 1A), in addition to the conventional augmentation pipeline, we employed an approach to train the cell image segmentation model using a modified CycleGAN process [22]. In comparison with conventional augmentation techniques, CycleGAN enhances the diversity of training cell images for the segmentation model by generating synthetic images of cells and their corresponding masks that are nearly indistinguishable from real images. The CycleGAN architecture is unique in that every step in the training process encompasses two mapping paths: forward consistency and backward consistency (Equation 1). This dual-path mechanism has the potential to introduce a greater degree of diversity to the training data. Within this architecture, the generator plays a pivotal role. It is capable of creating new synthetic cell images that might exhibit a significantly different distribution pattern from the original, real training samples. As a result, our segmentation model can train on a blend of these artificially generated images and the augmented real images. This hybrid training approach not only diversifies the data pool but also helps the model adapt to a broader spectrum of cell images. The addition of synthetic images simulates a wider array of scenarios that the model may encounter, thus improving its robustness and predictive power when faced with unfamiliar data.

For the synthetic image generator, we leveraged a 2D-UNET baseline architecture [23], as depicted in Figure S2. To further enhance the potential of the model for augmentation, we integrated a style-injecting network into this architecture [24]. This unique addition allows the infusion of style elements into the synthetic image generator to increase variation as well helping to mimic the natural diversity found in real microscopy cell images. This includes variations in cell shapes, structures, texture, and other subtle details that might influence the segmentation model’s accuracy. Our primary hypothesis is that our image generator architecture can incorporate more unique synthetic images into the training process to improve the segmentation model’s performance and robustness across different microscopy imaging styles and conditions.

We also employed two associated adversarial discriminators (D1 and D2, as seen in Figure 1A) within our framework [25]. The role of these discriminators is to incentivize the models to transform real images into synthetic counterparts that are virtually indistinguishable from the images within the real domain distribution [26]. To implement this, we adopted a PatchGAN discriminator-based architecture [27], to which we introduced a modification: a linear attention layer (Figure S3). The incorporation of linear attention, a strategy commonly used to enhance the performance of deep learning models [28], serves to enhance the performance of the PatchGAN discriminator. It accomplishes this by directing the discriminator’s focus toward the most pertinent features within the image. This focus ensures that the discriminator doesn’t get distracted by less meaningful image features, therefore improving its ability to differentiate between real and synthetic images. Consequently, this leads to the generation of superior-quality synthetic images that more closely mirror the real samples. Additionally, directing the discriminator’s attention to important features of an image contributes to the stability of the generator’s training process. This stability is vital, as it helps the generator produce consistently high-quality synthetic images over time, increasing the overall effectiveness of our CycleGAN-based training approach.

We also incorporated a differentiable image augmentation technique [29] into the input data for discriminator D1 during the training phase, an approach that has proven valuable (Figure S1B). Differentiable augmentation, in essence, applies identical random augmentations to both real and synthetic samples in a manner that is differentiable concerning the model parameters. This strategy motivates the discriminator less memorize the exact training samples and is effective in mitigating overfitting and boosting training stability, features that are particularly advantageous for GANs operating with limited data, thereby enhancing the overall performance of the image generator.

In the training process, we used a combination of generation and discrimination loss functions, where each loss function is assigned a specific weight to balance their contributions during optimization (Equations 3-14). We replaced the traditional L1 loss function [30] for the generator with a perceptual loss function, specifically using a pre-trained VGG network [31]. The perceptual loss function measures the high-level feature differences between the generated and target images, leading to a significant improvement in the quality of the synthetic images. The segmentation model employs Cross-Entropy and Dice losses, ideal for multi-class classification and imbalanced datasets, respectively [32]. Discriminators use Mean Squared Error (MSE) loss to differentiate between real and fake images [30]. By balancing the contribution of each loss function, the model ensures the production of high-quality synthetic cell images and accurate segmentation.

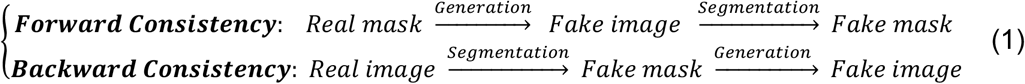

### Model’s performance evaluation

Because the main goal of this project was improving the segmentation model performance, we focused on segmentation precisions to evaluate the performance of our proposed architecture. In the testing phase, we assessed the performance of the segmentation model in three training scenarios by considering only the predicted cell body masks that have an Intersection Over Union (IOU) greater than or equal to the specified threshold. The precision at each threshold is then calculated as the ratio of true positives detected in cell bodies to the sum of true positives and false positives as commonly used for segmentation tasks (Equation 2) [12, 13]. In all experiments and score reports, we applied the cross-validation technique aiming to provide a more robust assessment of the segmentation model’s performance by reducing the impact of random variations in the training and validation data splits and ensuring that the model’s performance is not overly influenced by a specific subset of the data.

Figure 2A shows the conventional segmentation precision achieved with different training data sizes (introduced in Figures 1B-C) across different augmentation probabilities for two different cell types of our cell image dataset. As shown, initially, increasing augmentation could enhance accuracy by adding beneficial diversity to the training data. However, past a certain threshold (both cell types at p_vanilla=0.2), excessive augmentation might have introduced noise or irrelevant variability, potentially degrading model performance. We believe that the exact dynamics depend on the specific dataset, model, and augmentation techniques used, underscoring the need for task-specific experimentation and validation. Also, in our experiments, the impact of data augmentation varied between two cell-type images. The dataset comprising less complexity and lower diversity, Bronchial Epithelial Cells, showed a lower improvement from data augmentation compared to the more complex dataset of stem cell images. This can be attributed to the inherent simplicity of the data, which likely enabled the model to learn necessary patterns without the need for additional augmented examples. Conversely, for complex datasets with more inherent variability, data augmentation proved more beneficial by providing diverse examples, thus enhancing the model’s ability to generalize.

Next, we evaluated the performance of the segmentation model using the proposed CycleGAN-based training approach (with optimal values of p_vanilla=0.2 and p_diff=0.2). The proposed CycleGAN-based training approach can significantly improve the performance of the segmentation model at all IOU threshold values compared to when we train the segmentation model with the same limited dataset (AP=0.82 vs. AP=0.71 at IOU_thr=0.5 for stem cells and AP=0.87 vs. AP=0.83 for bronchial epithelial cells). Importantly, for the stem cells, the proposed approach even surpasses the performance of the model trained with 100% of the data. The CycleGAN optimal p_vanilla and p_diff parameters were derived from an extensive series of experiments in which various combinations of p_vanilla and p_diff were tested in incremental steps within the range of 0 to 1, which enabled us to locate the most effective values corresponding to the best validation scores.

We sought to test the impact of the modifications we have made to our CycleGAN-based model on the generation of synthetic images of cells (Figure 3). The figure compares synthetic cell images generated by three different model configurations: UNET with L1 loss, StyleUNET with L1 loss, and StyleUNET with VGG loss. From the figure, it is evident that the synthetic images produced by the UNET model with L1 loss show less detailed representation, achieving an average Fréchet Inception Distance (FID) score of 98. This score indicates a high dissimilarity between the synthetic and real cell images. The use of StyleUNET with L1 loss results in substantial improvements in image quality. The FID score drops to 43, signaling a closer match to the real cell images. This improvement primarily stems from the enhanced capability of StyleUNET to capture and generate variations in Style. Lastly, when we apply StyleUNET with the VGG perceptual loss, the synthetic images achieve a remarkably enhanced FID score of 23, reflecting a substantial increase in similarity to real cell images. In addition to the clearer depiction of the cell body, the model excels in generating high-quality representations of subcellular features such as the nucleus (Figure 4). This advancement signifies the superiority of the perceptual loss function in preserving high-level details and morphological nuances, thereby leading to more realistic synthetic images.

**Figure 4.**
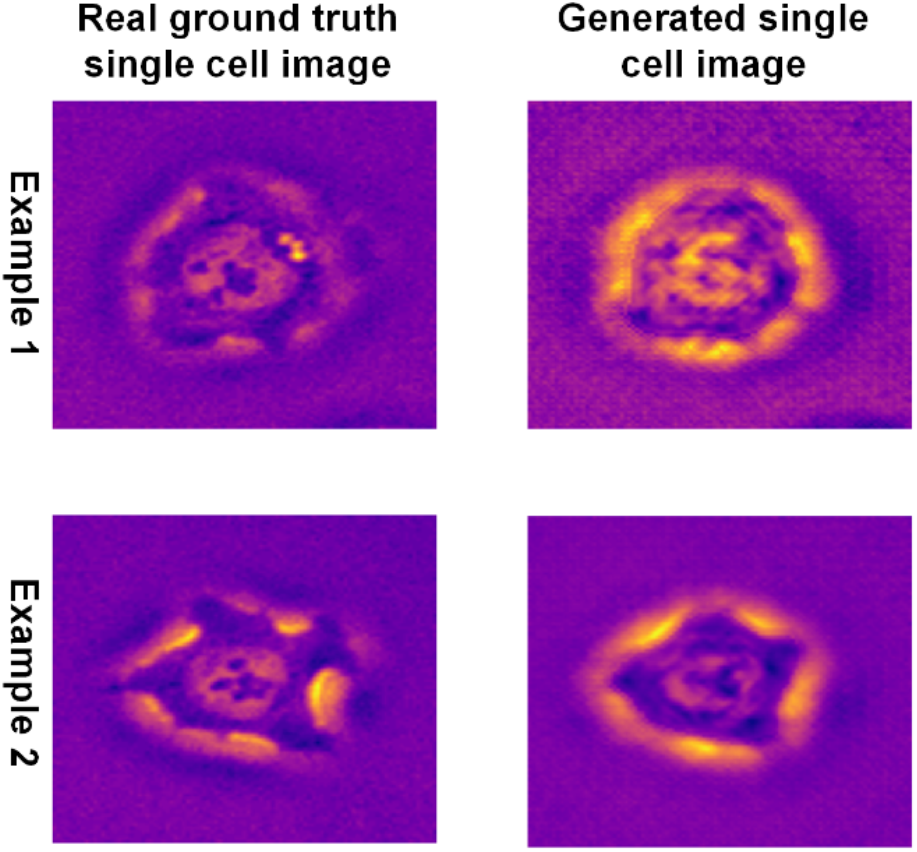
Two examples of pixel heatmap representation comparing the real with the synthetically generated cells by our modified CycleGAN network.

Most of our training data consisted of low and mid-density cell images. We were curious to test if our generator is able to extrapolate its knowledge by generating new synthetic high-density images of cells that were not seen during training. This is particularly useful as manual annotation of high-density images of cells can be very time-consuming and error-prone. To accomplish this, we designed an algorithm for generating synthetic high-density cell masks (a relatively easy task) as input for the generator (in the test phase). As shown in Figure 5, our approach confirms the ability to extrapolate knowledge from low and mid-density cell images, creating annotated images across any density level and magnification, even those absent in the original dataset. Such a capacity enables us to create synthetic cell images mimicking a broad range of real-world scenarios. The capacity of our model to extrapolate learned knowledge to unseen scenarios can provide a powerful tool to generalize this approach aiming to develop the segmentation models for a variety of cellular imaging modalities.

**Figure 5.**
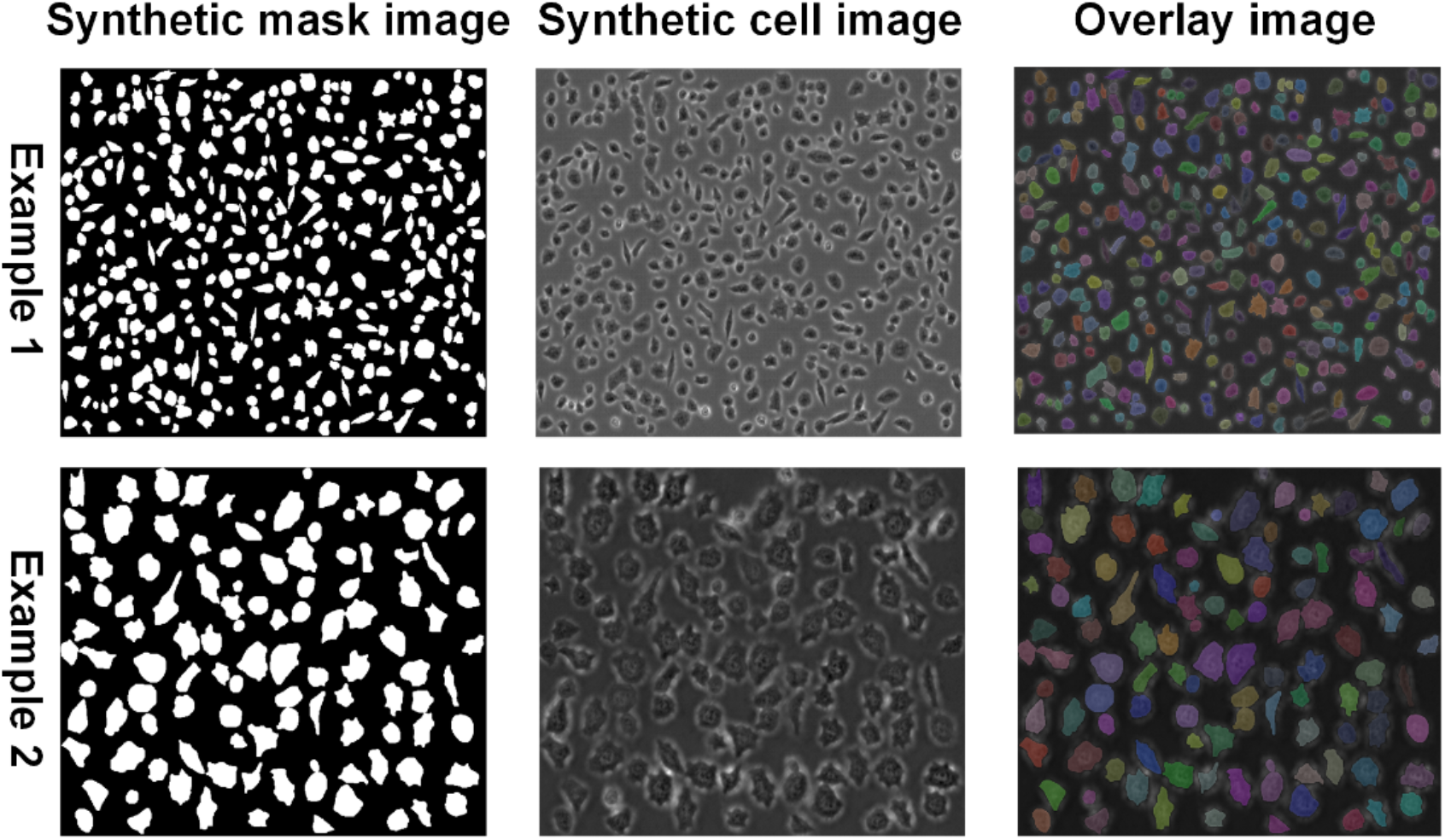
CycleGAN method is able to generate unseen imaging scenarios. Two examples of synthesis of unseen high-density cell images with different magnifications using the trained StyleNet generator.

## Discussion

A large and diverse annotated dataset of images is key to the successful development of deep-learning models that can perform across a variety of real-world images. Currently, a harmonized large and diverse dataset of microscopy images is not available to train new deep-learning models mostly because annotation of many microscopy images is a tedious and time-consuming task. Our study provides a solution to this problem by proposing a novel method for training cell segmentation models using a CycleGAN approach to address the critical issue of limited annotated data in cellular imaging. By harnessing the potential of GANs, we effectively generated a diverse set of synthetic realistic cell images, enhancing the diversity of the available training datasets without manual annotation and improving the overall performance of the segmentation models with a limited annotated dataset. Importantly, we showed that the CycleGAN approach allows for the extrapolation of knowledge by model by generating synthetic images that the model has not been exposed to during the training.

We made several modifications to the original CycleGAN architecture to be able to apply to the microscopy images. First, a style generation path was integrated into the synthetic image generator to boost variation in synthetic images. Second, a linear attention mechanism was incorporated into the PatchGAN discriminator-based architecture to enhance its differentiation capabilities and synthetic image quality. Third, differentiable image augmentation was introduced during the training phase to further diversify image generation and reduce the risk of overfitting. Fourth, instead of the L1 loss function conventionally used in the CycleGAN, we employed a combination of Cross-Entropy (CE) and Dice losses for the segmentation, improving the handling of multi-class classification and imbalanced datasets. Finally, as a critical modification, we replaced the L1 loss function in the generator with a VGG perceptual loss function to promote the retention of more high-level features and nuances in the generated synthetic images, leading to enhanced similarity between real images of cells and synthetic images. These enhancements collectively improved the diversity and quality of synthetic cell images, resulting in a more diverse and generalized segmentation model trained with various microscopy imaging styles and conditions.

Our experimental results show that the proposed CycleGAN approach provides a straightforward solution for the paucity of annotated microscopy data for training deep-leaning models. Experimental results showed that the performance of segmentation models trained using our CycleGAN-based method improved the segmentation precision across two cell types of our dataset (11% precision improvement in stem cells and 4% improvement in Bronchial epithelial cells). Notably, this enhancement was observed irrespective of the scarcity of annotated cell image datasets, illustrating the potential of our approach in effectively addressing this prevalent issue in biomedical imaging. Implementation of Style injecting in our UNET generator significantly improved the quality of synthetic images, reflected by an FID score reduction from 98 to 43. A further enhancement was achieved by substituting the conventional L1 loss function with a VGG perceptual loss, resulting in an FID score of 23.

In conclusion, our CycleGAN-based method opens up new possibilities for training deep-learning models for microscopy applications by offering a novel solution to the challenge of limited annotated cell image datasets. This study illustrated how generative deep learning methods like GANs can be utilized to address data limitations in microscopy, thereby pushing the boundaries of what’s possible in the field of biomedical imaging. It is important to mention that while our approach has shown promising results, there is room for further improvement and experimentation, including exploring different GAN architectures and further refinement of the augmentation techniques. The generated synthetic images can also be made more diverse and realistic through additional modifications in the GAN training process.

### Methods Dataset

For the training and validation of our deep learning models, we utilized the DeepSea dataset [12]. This dataset comprises a collection of time-lapse microscopy images of cells, along with manually annotated cell body masks, making it a good resource for applications like cell segmentation and tracking. It consists of three types of cell images: 1) Mouse Embryonic Stem Cells (1074 images), 2) Bronchial epithelial cells (2010 images), and Mouse C2C12 Muscle Progenitor Cells (540 images). We only used Embryonic Stem Cells and Bronchial epithelial cells due to their adequate dataset sizes. We used 15% of the dataset samples for testing all training scenarios. For the first (Figure 1A) and second scenarios (Figure 1B), we randomly selected subsets representing limited datasets, 25% of the remaining samples. Also, all the remaining samples (excluding the 15% test samples) were employed as the training set for the third scenario (Figure 1C).

### Segmentation model

We used the DeepSea segmentation model [12] in our experiments, and it can be replaced by any other segmentation model, depending on the segmentation application. It is shown that the DeepSea model can segment the DeepSea dataset samples well. It is an efficient scaled-down version of the 2D-UNET model. To simplify the proposed idea, we chose not to incorporate the layer representing touching cells, as this would necessitate custom touching cell masks (alongside cell body masks) and additional loss functions.

### Augmentation functions

Image augmentation techniques play a critical role in expanding the diversity of training datasets, thereby improving model generalization and robustness [33, 34]. In the training process of our deep learning models, we applied some mostly used conventional image augmentation functions to every single cell image with the probability of p_vanilla, including random histogram equalization, random crop, random sharpness adjustment, random brightness adjustment, random contrast adjustment, random horizontal flip, random vertical flip, random saturation, adding random gaussian noise, and adding random gaussian blur as shown in Figure S1A. For the binary mask images, we only used the applicable random crop, horizontal flip, and vertical flip functions. The training algorithm executes a sequence of the provided augmentation functions for each cell and mask image pair with a pre-defined probability value ‘p_vanilla’. In the requested augmentation pipeline, each function is randomly chosen with a consistent probability of 50% and is also applied in a randomized sequence.

However, when it comes to training Generative Adversarial Networks (GANs), especially models like CycleGAN that learn mappings between different image domains, conventional augmentation might not be sufficient for enhancing the diversity of generated images. This is where differentiable augmentation, as proposed in [29], becomes valuable. Differentiable augmentation applies the same random augmentations to both real and fake samples in a way that is differentiable with respect to the model parameters. This approach encourages the discriminator to less memorize the exact training samples, thus causing the generator to produce more diverse images, thereby improving the overall image generation performance. Furthermore, differentiable augmentation can mitigate overfitting and improve training stability, making it particularly beneficial for GANs trained with limited data. In this project, we used five different differentiable augmentation functions such as random contrast, random brightness, random cutout, random translation, and random saturation. The decision to perform an augmentation is dictated by the probability variable ‘p_diff’. In an attempt to ensure fair representation and randomness, each of the differentiable augmentation functions is executed in a randomized sequence, with each having an equal 50% probability of selection. This approach not only diversifies the images but also ensures that the model remains adaptable to any new form of data it might encounter in the future, thus improving its resilience and overall effectiveness.

**Figure S1.**
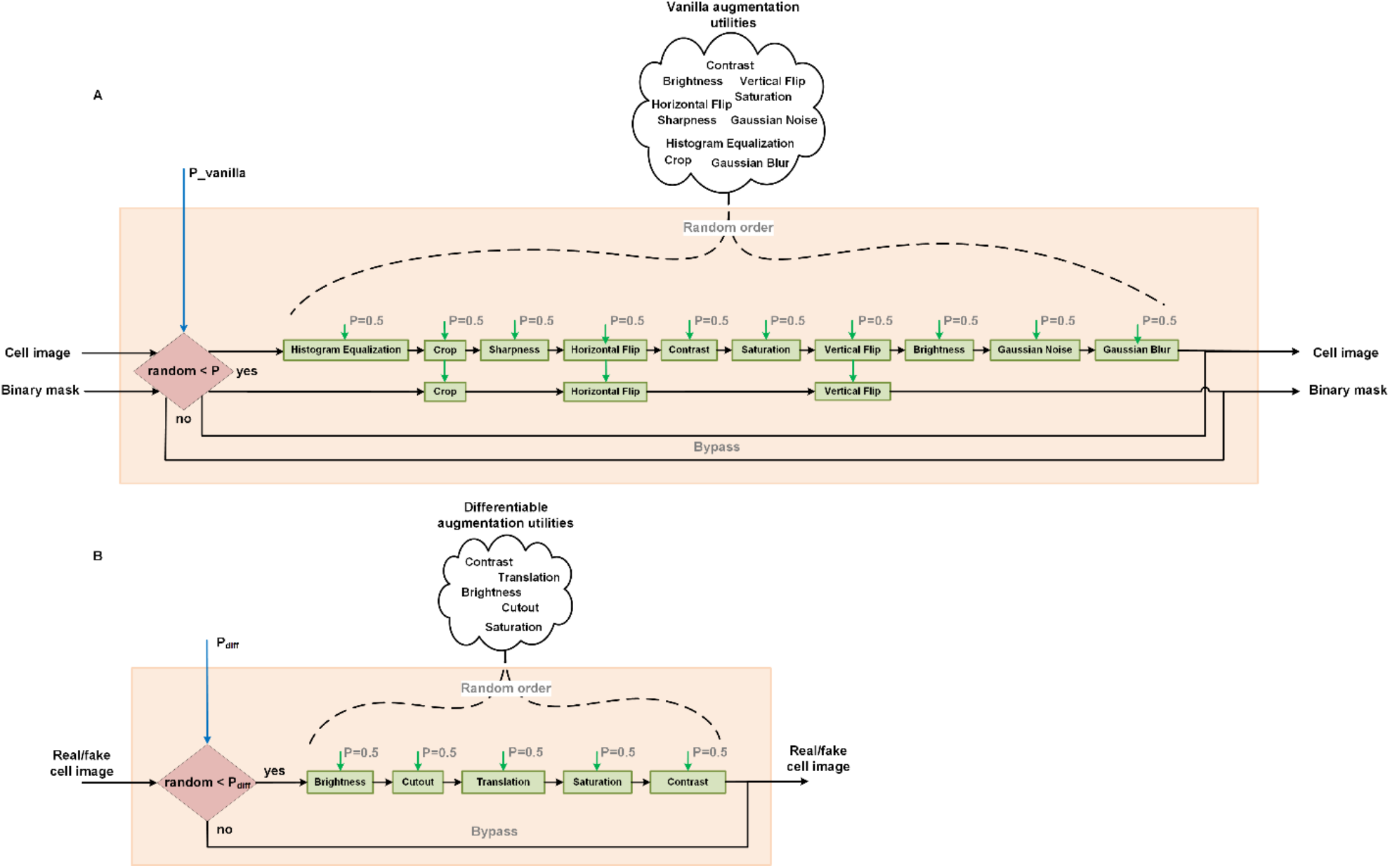
Augmentation functions used in the proposed training process. A) The conventional vanilla augmentation that was applied to both cell image and corresponding binary mask pairs aimed to increase the diversity of samples fed into the CycleGan-based model. B) The differentiable augmentation functions used to reduce the risk of overfitting and help the generator in producing more diverse synthetic images.

### Generator model

In our segmentation training approach, the generator is responsible for generating synthetic cell images. It employs a 2D-UNET architecture [23], as shown in Figure 2S. UNET is renowned for its effectiveness in biomedical image segmentation due to its unique architecture, which consists of a contracting path to capture context and a symmetric expanding path that enables precise localization. However, we have taken this a step further by incorporating a style decoding path into the decoder part of the UNET architecture, an idea inspired by the StyleGAN2 model [24].

This fusion of concepts from StyleGAN2 and UNET brings about the prospect of generating better synthetic images. The style decoding network is designed to control the stylistic aspects of the generated images, thereby allowing the model to create more diverse and potentially higher-quality synthetic cell images. This combination of architectures seeks to maximize the strengths of both models - the segmentation prowess of UNET and the sophisticated generative capacity of StyleGAN2. This integration could potentially yield a more powerful generator model for synthetic cell image creation, thereby enhancing the overall performance of our CycleGAN-based cell image segmentation system.

**Figure S2.**
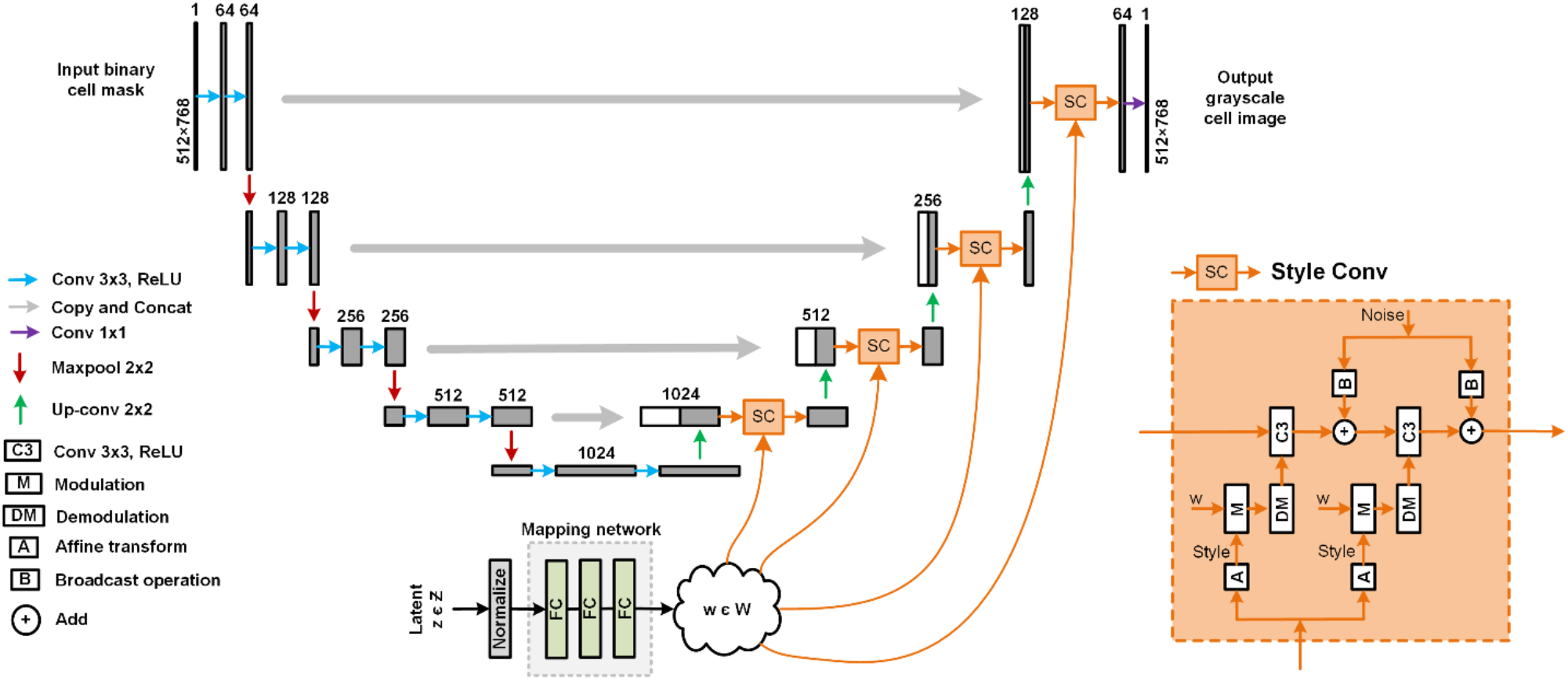
Generator architecture. It employs the 2D-UNET architecture with a style decoding network to create more diverse and potentially higher-quality synthetic cell images.

### Discriminators

In our proposed method, we applied a modified version of the PathGAN baseline architecture [27] for our discriminators, integrating a layer of residual linear attention, as shown in Figure S3. The PathGAN architecture, known for its effectiveness in examining both global and local image features, has demonstrated impressive performance in diverse GAN applications. The architecture operates by creating multiple ‘paths’ with different receptive field sizes, enabling the model to scrutinize image details at various scales.

However, we sought to improve the discriminator’s ability to focus on critical features by incorporating an additional layer of residual linear attention [28]. This is an approach to attention mechanisms that makes use of a linear combination of input features and learned attention maps, thereby enabling the model to weigh different regions of the input differently. As a result, the model can focus on more critical parts of the image, thereby enhancing its ability to discriminate real images from synthetic ones accurately. By modifying the PathGAN discriminator, we aimed to improve the model’s focus on salient image features, thus boosting its ability to accurately distinguish between real and generated images.

**Figure S3.**
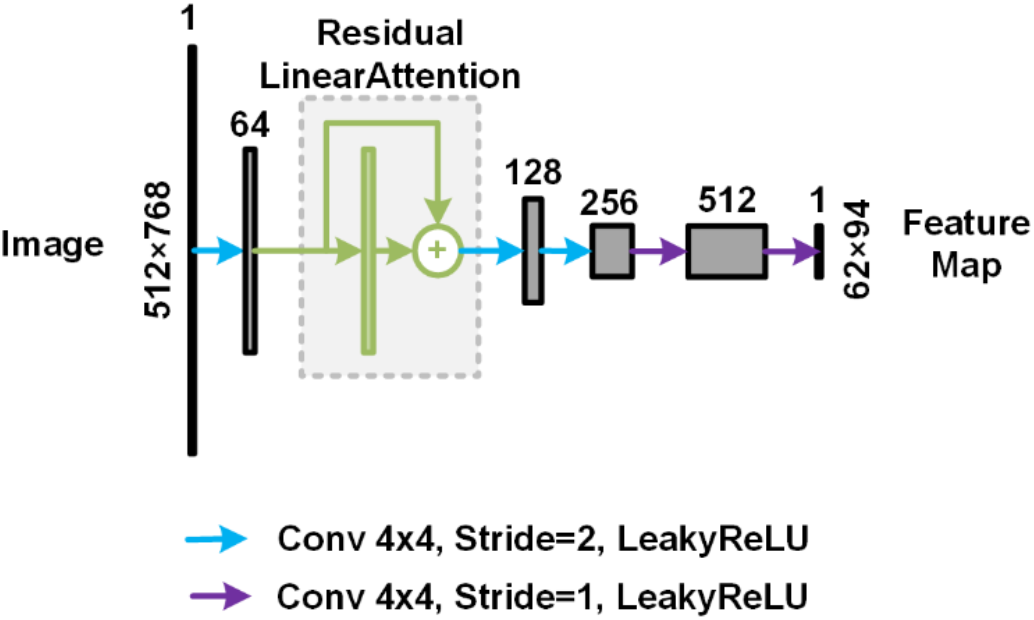
A modified version of the PathGan discriminator used in our proposed approach. We integrated a layer of residual linear attention to improve the discriminator performance.

### Evaluation metrics

In the testing phase, we leveraged the Intersection over Union (IoU) metric, also known as the Jaccard Index, which ranges from 0 to 1, to evaluate the alignment between the segmentation model’s predictions and the manually annotated ground truth masks [35]. For each test image, we designated each detected cell body as a True Positive (TP) prediction if its IoU index exceeded a predetermined threshold value, indicating a valid match to the ground truth. Conversely, any ground truth cell body masks that failed to find a valid match were classified as False Negatives (FN), and any predictions lacking corresponding ground truth masks were labeled as False Positives (FP), representing non-cell entities. Subsequently, we calculated the Average Precision (AP) value for each image in the test set using Equation (2). AP, frequently employed by state-of-the-art methods for cell body segmentation tasks, provides a single-figure measure of quality across recall levels [12, 13].

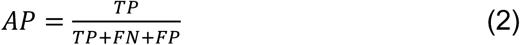

Frechet Inception Distance (FID) is also a widely used metric to evaluate the quality of images generated by GAN models [37]. It quantifies the dissimilarity between the distributions of generated and real images in the feature space of a pre-trained Inception network. Lower FID scores represent higher-quality synthetic images that more closely resemble the distribution of real images. By assessing differences in both mean and covariance of the features, FID provides a more comprehensive evaluation of image quality and diversity, making it a good choice for evaluating the performance of our modified generator.

### Loss functions

In the training phase of our CycleGAN-based model, we employed a series of loss functions to effectively optimize the performance of both the generator and the segmentation model. These loss functions are specifically tailored to address the unique challenges presented by the task of generating high-quality synthetic images and accurately segmenting cell structures.

Two fundamental loss functions utilized in our model are identity loss and reconstruction loss. Identity loss ensures that an image translated to its own domain remains unchanged, which encourages the generator to preserve color and texture composition between the input and output [22]. Reconstruction loss, on the other hand, is used to maintain cycle consistency, ensuring that an image translated from one domain to another can be accurately translated back to its original form. These losses are critical to ensure that the model not only learns the correct mappings between the domains but also produces images that are consistent with the original data distribution.

For the generator, we utilized the VGG perceptual feature loss function [31, 36] for both identity and reconstruction loss (Equations 5 and 7). The VGG loss function is a high-level feature extraction loss that helps in preserving the perceptual and semantic understanding of the images. It is a concept based on a deep convolutional neural network (CNN), like VGG, that has been pre-trained on a large dataset for an image classification task.

For the segmentation model, we combined Cross-Entropy (CE) loss and Dice loss for both identity and reconstruction loss (Equations 6 and 8) [32]. Cross-Entropy loss is a popular choice for multi-class classification problems, calculating the dissimilarity between the predicted probability distribution and the ground truth distribution. The Dice loss, on the other hand, is specifically designed for handling imbalanced datasets and is extensively used in medical image segmentation tasks due to its efficiency in dealing with small objects and imbalanced classes. By using these two loss functions in tandem, we enhance the performance of our segmentation model, ensuring it can effectively handle the challenges of cell image segmentation.

The discriminators in our model were optimized using the Mean Squared Error (MSE) loss as an adversarial loss. This loss function encourages the discriminators to distinguish between real and fake images by minimizing the average squared differences between the predicted and actual values.

Each of these loss functions is assigned a specific weight in order to balance their contributions during the optimization process (Equations 13 and 14). By integrating these diverse loss functions and carefully selecting their weights, we can effectively train our CycleGAN-based model, ensuring both the production of diverse, high-quality synthetic cell images and the accurate segmentation of cell structures.

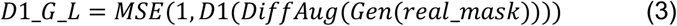

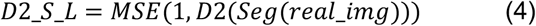

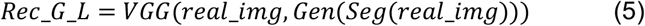

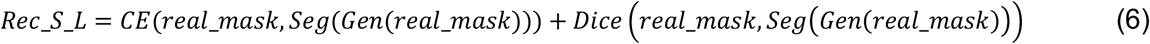

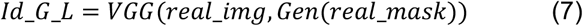

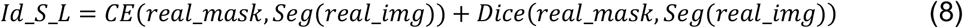

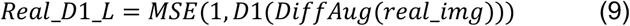

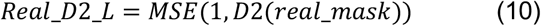

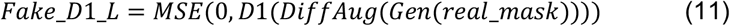

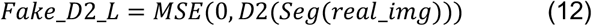

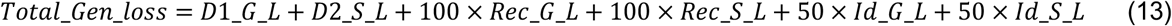

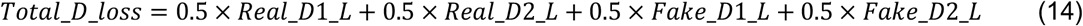

## Code and data availability

The methods we’ve implemented are available as Python scripts and can be downloaded from https://github.com/abzargar/GANSeg. The DeepSea dataset of images can be downloaded from https://deepseas.org/datasets/

## Author Contribution

A.Z.K and S.A.S conceived the project and wrote the manuscript. A.Z.K and N.M also implemented the related Python scripts and prepared the manuscript materials.

